# Parallel evolution of chimeric genes in microbial evolution experiments

**DOI:** 10.64898/2026.06.14.732165

**Authors:** Rohan Maddamsetti, Mehmet Orman

## Abstract

Recombination can generate new or improved proteins by merging pieces of pre-existing genes into a new whole. Here, when we reanalyzed genomic data from several laboratory evolution experiments with *Escherichia coli*, evidence for the convergent evolution of chimeric genes became apparent. In these experiments, a pair of paralogous genes were recombined into a single hybrid copy by large genomic deletions. In one case, the excision of a cryptic e14 prophage occurred in 6 out of 8 replicates of a 22-day fluoroquinolone-resistance evolution experiment. These parallel prophage excisions recombined an *icd* isocitrate dehydrogenase gene with a homologous *icdC* C-terminal fragment pseudogene. In the other case, convergent 23 kB deletions recombined *cpsG* paralogs across replicate populations of the Lenski long-term evolution experiment with *Escherichia coli* (LTEE), generating a chimeric phosphomannomutase. Together, these results indicate that chimeric genes evolve rapidly in bacteria and are more common than previously believed, because they are easily missed by standard genomic analyses. Experiments are needed to determine whether these chimeric proteins are adaptive, are by-products of adaptive genomic deletions, or both.

**Data Summary:** All data used in this study are publicly available. All data, code, and results for this study are available on Zenodo (DOI: 10.5281/zenodo.20692138). Genomic data published by Mohiuddin et al. are available in the European Nucleotide Archive (Study ERP166963 with Study Accession PRJEB83321), and genomic data published by Tenaillon et al. are available from the NCBI BioProject database under accession number PRJNA294072.

**Impact Statement:** Genomic deletions can generate chimeric proteins by splicing together parts of existing genes. However, the extent to which chimeric proteins form, and the extent to which they are adaptive are not well-understood. Complex structural variants (such as chimeric protein formation) are often missed or incompletely annotated by standard genomic pipelines. Furthermore, there is an ongoing need for new methods to detect selection on complex structural variants. In this work, we report evidence of chimeric gene formation in laboratory evolution experiments with *Escherichia coli*. In both experiments, chimeric genes evolved in parallel across independent replicate populations, which is an unambiguous signal of positive selection (although it remains unclear whether selection is operating on the chimeric gene, the genomic deletion creating the chimeric gene, or both). The observation of chimeric gene formation in laboratory experiments indicate that chimeric genes evolve rapidly in bacteria and are more common than previously believed. Therefore, this work points to an important new direction for microbial genomics research: what is the extent of chimeric gene formation in natural bacterial populations, and what consequences might chimeric genes have for bacterial adaptation?

## Introduction

New genes and functions often evolve by duplicating, inserting, deleting, and recombining parts of existing genes, leading to functional innovation^1-3^. Due to advances in genome sequencing technology^4^, it has become clear that such structural variation is pervasive across the tree of life^5^. The potential for recombination to generate new genes has long been appreciated in eukaryotes^6,7^. It has recently become clear that recombination and structural variation are also pervasive in natural bacterial populations^8^ and during microbiome evolution^9^. For this reason, the extent to which duplications, deletions, and recombination generate adaptive protein variation in bacteria remains poorly understood.

Genomic deletions are usually considered as a mechanism that streamlines genomes by removing unneeded genes. However, interstitial genomic deletions can in fact yield new genes. There are two primary mechanisms. First, a deletion can yoke together parts of unrelated genes, forming a new protein by gene fusion^10,11^. Second, a deletion can yoke together different parts of homologous genes (such as paralogs formed by recent gene duplications), forming a new chimeric gene. In this paper, we describe two cases of chimeric gene formation by this second mechanism across different evolution experiments with *Escherichia coli*. These findings suggest chimeric gene formation may be more common than previously believed, because many more cases of chimeric gene formation in bacteria may have gone undetected in existing genomic studies of bacterial evolution in the laboratory and in nature.

## Results

### Parallel evolution of a chimeric isocitrate dehydrogenase during experimental evolution under fluoroquinolone exposure and recovery

Mohiuddin et al. carried out an evolution experiment with *Escherichia coli* K-12 to investigate adaptation to fluoroquinolone stress. In this 22-day experiment, 8 replicate populations were subjected to multiple cycles of high-dose ofloxacin exposure followed by drug-free recovery^12^. Whole-genome sequencing of each evolved population revealed limited evidence of convergent evolution. This result was surprising, because parallel evolution is a more typical outcome for short-term evolution experiments with large population sizes and strong selection pressures such as antibiotic use^13^. Since parallel evolution is mediated by mutations with either large fitness benefits or high rates^14,15^, we asked whether some kinds of mutations (such as gene copy number changes) may have been missed by the initial analysis, which focused on single nucleotide polymorphisms (SNPs), insertions and deletions (indels) and structural variants (SVs).

To answer this question, we re-analyzed these genomic data using *breseq*^16-18^ and additional statistical methods to detect copy number variation^13,19^. While we did find evidence of parallel 2-3× gene copy number amplifications of the *bamA, rpoB*, and *lldP* genes (Supplementary Table S1), we further examined SNPs in the *icd* isocitrate dehydrogenase gene, because *icd* has been implicated in persistence and antibiotic tolerance^20^ and has been reported to be recurrently mutated in genomic studies of tolerant populations^21^. 6 out of the 8 evolved populations show multiple identical synonymous changes to *icd*, including mutations affecting the same codon. In addition, these synonymous changes correlated perfectly with missing sequencing coverage affecting the C-terminus of *icd*, indicating parallel 15.1 kilobase-pair (Kbp) deletions (Table 1). By examining the K-12 MG1655 genome in the Benchling^22^ and Proksee^23^ genome browsers, we found that these putative deletions correlated perfectly with the coordinates of cryptic prophage e14 in in the MG1655 genome (Table 1). Notably, e14 is a known prophage element in *E. coli* K-12 that has been shown to excise under induction of the SOS response^24-26^, and has been linked to *icd* mutations in ciprofloxacin-evolved populations^27^. Further visual inspection of the *icd* and prophage e14 locus in MG1655 revealed that cryptic prophage e14 is flanked by *icd* at one end, which extends into the e14 prophage, and by an *icdC* C-terminal pseudogene at the other end of the e14 prophage, consistent with observations reported by Hill et al.^26^ and Sulaiman et al.^27^ (Figure 1A). To further test if the parallel 15.1Kbp deletions of prophage e14 can recombine *icd* with *icdC*, we modeled the effect of deleting the region spanning 1196266-1211412 in the MG1655 genome on the *icd* gene and aligned the resulting chimeric protein against its parental genes *icd* and *icdC* (Figure 1B). The resulting protein has a duplicated MLRHMGWTEAADLIVKGME C-terminal sequence and D417E and D429E substitutions through recombination with *icdC*, but is otherwise identical to the translated protein product of *icd* (Figure 1B).

**Table 1.**
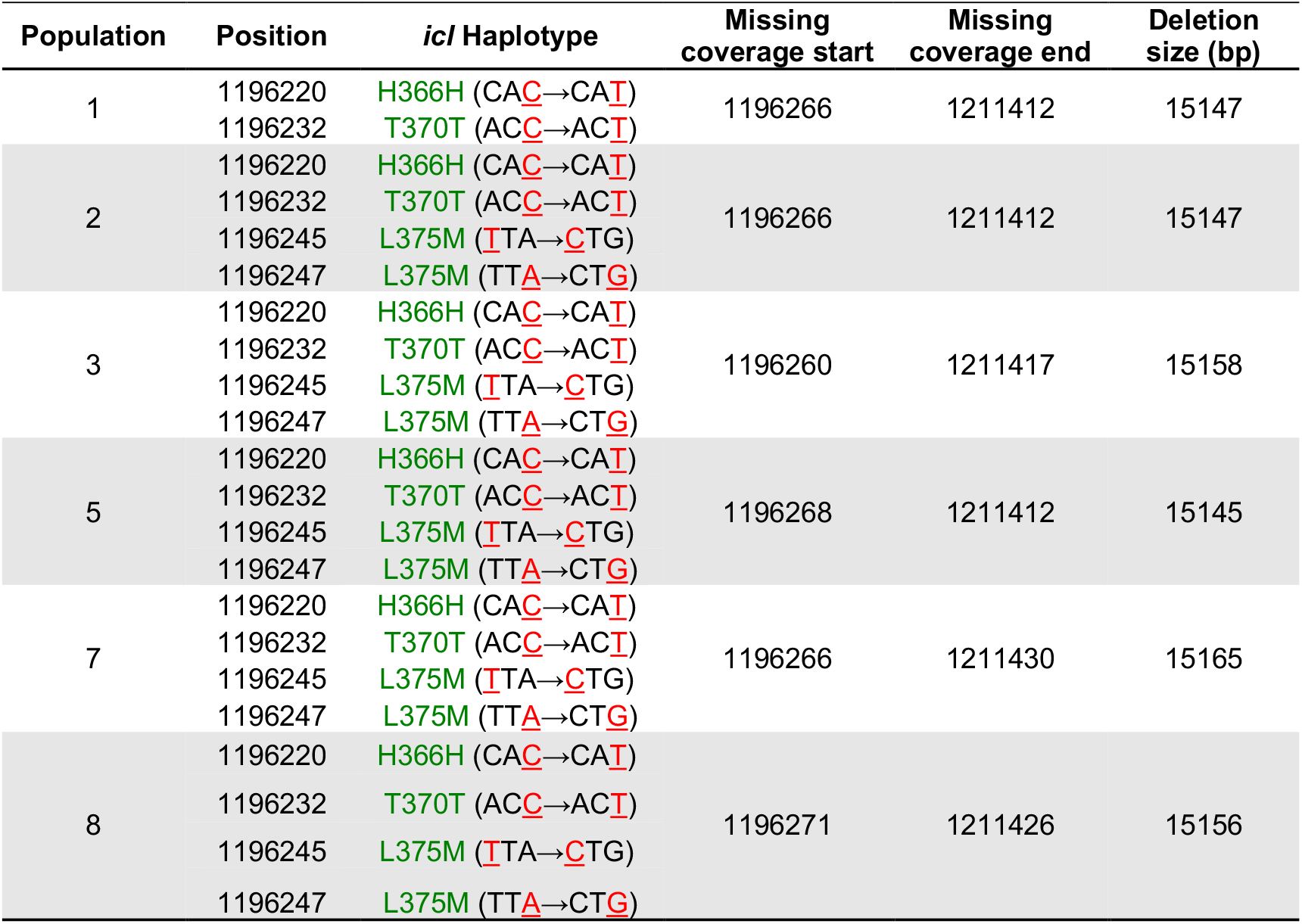
Parallel synonymous changes in *icl* isocitrate dehydrogenase correlate with loss of cryptic prophage e14 (1196209-1211412) during experimental evolution of fluroquinolone resistance (Mohiuddin et al. 2025).

**Figure 1.**
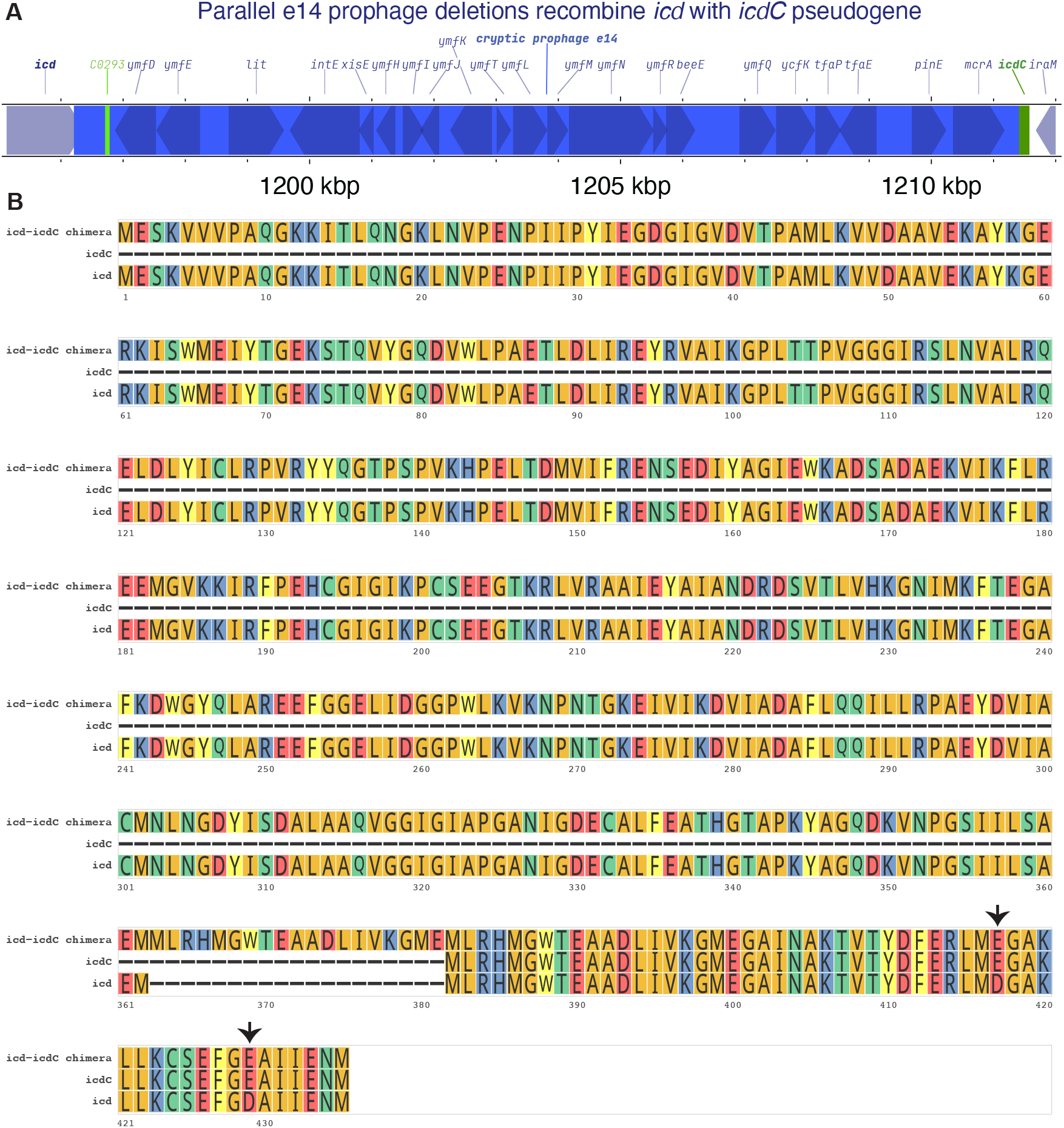
Parallel e14 prophage deletions recombine *icd* with *icdC* pseudogene during experimental evolution of fluoroquinolone resistance. A) Structure of pre-deletion locus in the *Escherichia coli* K-12 MG1655 ancestral clone. Protein-coding genes are colored gray-blue, a small RNA is colored light green, the *icdC* pseudogene is colored dark green, and e14 prophage is colored blue. B) Multiple protein sequence alignment of chimeric *icd*-*icdC* with its parental *icd* and *icdC* genes. Arrows indicate *icdC* amino-acid substitutions introduced by recombination.

### Parallel losses of the rfb O-antigen (O-polysaccharide) biosynthesis gene cluster drive evolution of chimeric phosphomannomutase in the LTEE

A similar phenomenon occurs in the Lenski long-term evolution experiment with *E. coli* (LTEE). Parallel 23 kbp genomic deletions occurred in 8 of the 12 replicate populations of the LTEE^28^ (Table 2). These 23 kbp deletions recombine two highly similar homologs of *cpsG*, which encodes a phosphomannomutase involved in the biosynthesis of the extracellular polysaccharide colanic acid^29^. Visual inspection of this region of the *E. coli* B str. REL606 genome in the Benchling^22^ and Proksee^23^ genome browsers shows that these deletions affect the *rfb* O-antigen (O-polysaccharide) biosynthesis gene cluster, although this strain already does not produce O-specific side-chain lipopolysaccaride^30^ (Figure 2A). We examined the effect of this Δ23,293 bp mutation on *cpsG* starting from position 2,031,703, as this variant occurs in two LTEE populations, Ara+4 and Ara–1 (Table 2). An alignment of the resulting chimeric protein shows that the deletion recombines the N-terminus of CpsG with the C-terminus of ManB (Figure 2B).

**Table 2.** Parallel genomic deletions between *manB* and *cpsG* phosphomannomutase paralogs in the LTEE*.

| Population | Fixation | Position | Mutation | Gene |
| --- | --- | --- | --- | --- |
| Ara+3 | Yes | 2,032,210 | Δ23,293 bp | *[manB]–[cpsG]* |
| Ara+4 | Yes | 2,031,703 | Δ23,293 bp | *[manB]–[cpsG]* |
| Ara–1 | Yes | 2,031,703 | Δ23,293 bp | *[manB]–[cpsG]* |
| Ara+5 | No | 2,032,369 | Δ23,293 bp | *[manB]–[cpsG]* |
| Ara–2 (L<sup>**</sup>) | Yes | 2,032,399 | Δ23,293 bp | *[manB]–[cpsG]* |
| Ara–2 (S<sup>**</sup>) | Yes | 2,031,537 | Δ23,293 bp | *[manB]–[cpsG]* |
| Ara–3 | Yes | 2,032,711 | Δ23,293 bp | *[manB]–[cpsG]* |
| Ara–4 | Yes | 2,032,114 | Δ23,293 bp | *[manB]–[cpsG]* |
| Ara–5 | Yes | 2,032,762 | Δ23,293 bp | *[manB]–[cpsG]* |

\*Data retrieved from https://barricklab.org/shiny/LTEE-Ecoli/ on May 4 2026.
\*\*Data about “large” and “small” colony-forming ecotypes in the Ara–2 population retrieved from Supplementary Data 1 of Tenaillon et al. (2016).

**Figure 2.**
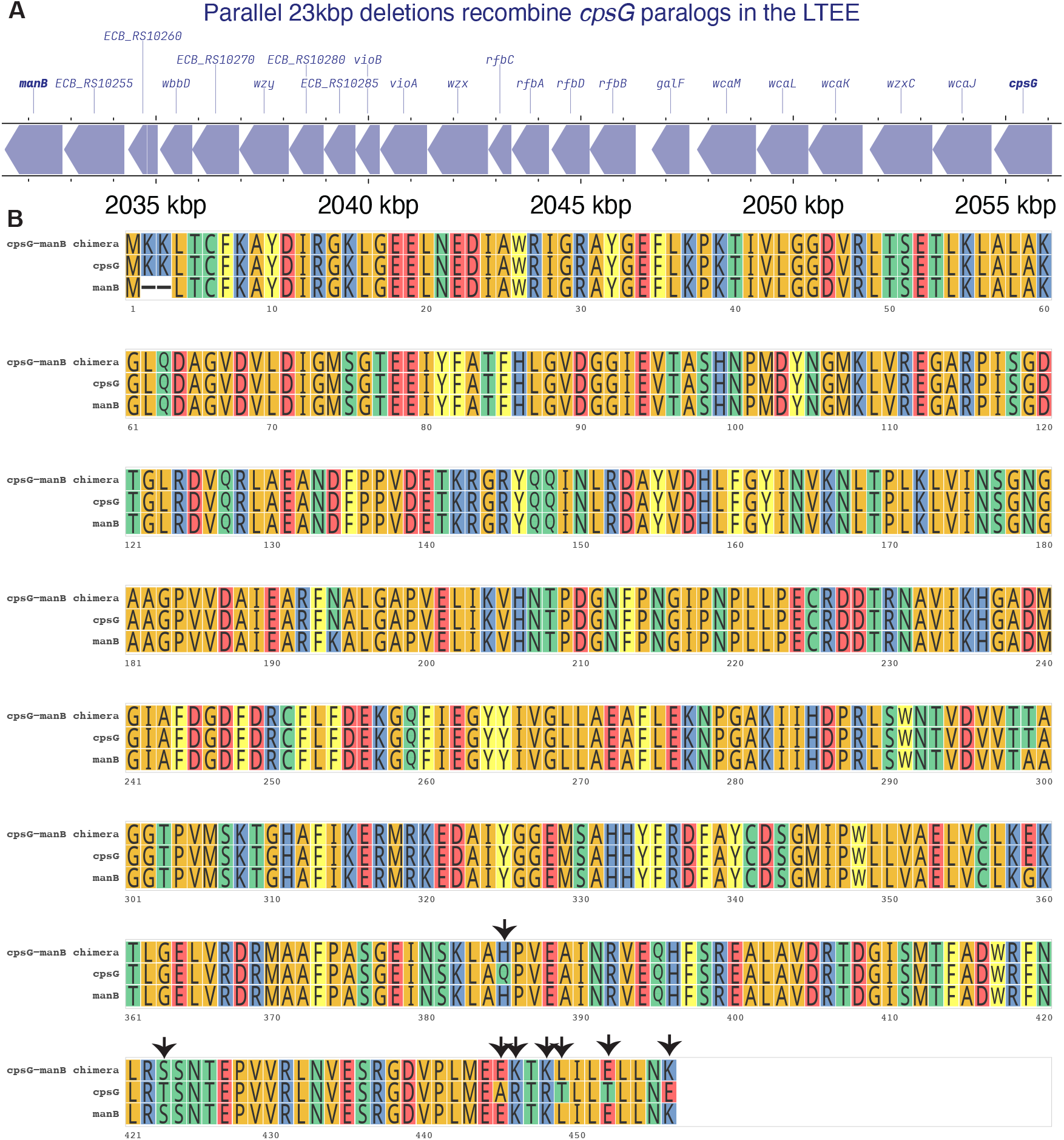
Parallel 23 kbp deletions recombine *cpsG* paralogs in the LTEE. A) Structure of pre-deletion locus in the genome of the REL606 LTEE ancestral clone. Protein-coding genes are colored gray-blue. B) Multiple sequence alignment chimeric *cpsG-manB* phosphomannomutase with its parental *cpsG* and *manB* genes. Arrows indicate *manB* amino-acid substitutions introduced by recombination.

## Discussion

Here, we describe the parallel evolution of chimeric proteins via large genomic deletions in laboratory evolution experiments. In both cases, the large deletion was flanked by two highly homologous gene sequences, likely derived by recent gene duplication. Recombination between those paralogs joined the N-terminus of one protein with the C-terminus of the other, removing all intervening genes (Figure 3).

**Figure 3.**
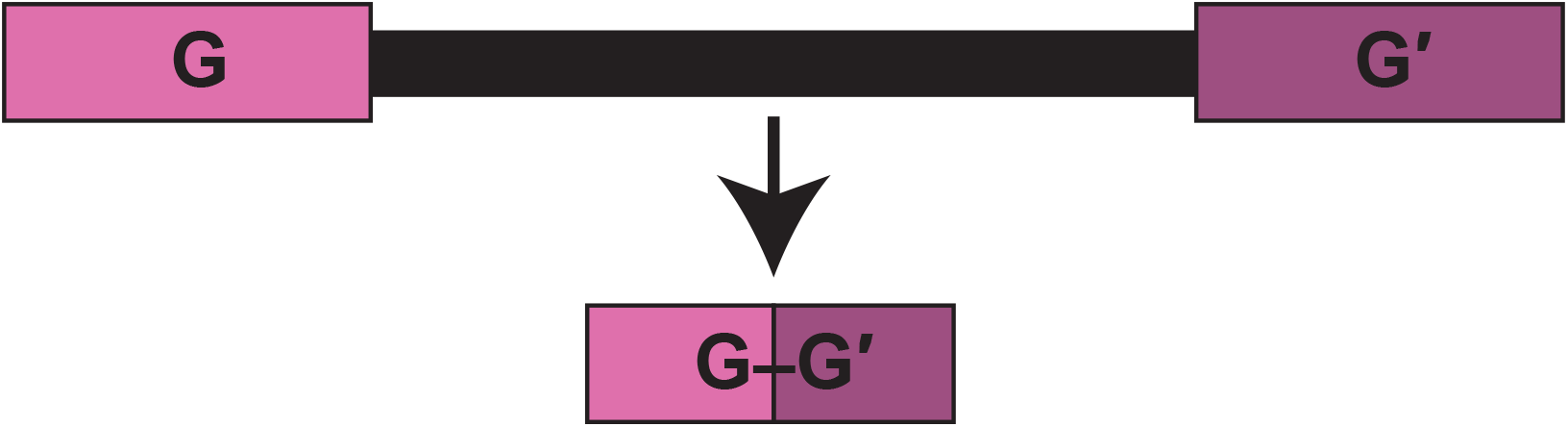
Large genomic deletions recombine paralogous genes G and G′, resulting in chimeric gene G–G′.

While aspects of these events were reported previously^12,27,28^, the evolution of chimeric isocitrate hydrogenase was not explicitly recognized by Mohiuddin et al. because two separate observations had to be combined to make this inference: an anomalous pattern of parallel synonymous substitutions, coupled with loss of sequencing coverage at a neighboring cryptic e14 prophage^12^. In the second case, it was known that the parallel 23kbp deletions affected *cpsG* homologs in the LTEE, but the resulting chimeric *cpsG* gene sequences had not been studied further or reported in any publications^28^.

We speculate that many more cases of chimeric gene evolution may have occurred but gone undetected in previous studies. Gold-standard microbial genome analysis pipelines like *breseq* report deletions as well as synonymous mutations introduced by recombination, but to our knowledge, no current genomic pipelines directly annotate new chimeric proteins that arise in evolution experiments. Furthermore, variation at recently duplicated genes remains largely uncharacterized in evolution experiments. For instance, the dynamics of genetic variation at the *manB* and *cpsG* in the LTEE are poorly understood, because large-scale genomic and metagenomic studies of the LTEE filtered out all genetic variation in repetitive regions, including multiple direct sequence repeats shared by *manB* and *cpsG*^28,31^.

This work leaves open the question of whether the parallel evolution was driven by positive selection on the chimeric gene, positive selection on the deletion, or both. The second explanation (selection on the deletion) is more parsimonious. Large genomic deletions can be beneficial because they remove unneeded proteins. They can also occur at high rates due to homology between direct genetic repeats, as found in paralogous gene copies. Genomic deletions could also occur at high rates due to the induction of site-specific recombination systems, such as those often present in some prophage^32^. This seems to be the case for e14 prophage excision and *icd* chimera formation, which are specifically induced by the SOS DNA damage response^24-26,32^. Nevertheless, the weight of current evidence suggests that positive selection is still a key driver for the parallel evolution of *icd* chimera formation in the experiments conducted by Mohiuddin et al.^12^ and by Sulaiman et al.^27^, because e14 is excised slowly and/or inefficiently after SOS induction^24,25^. Further experiments with isogenic *E. coli* K-12 strains with and without excised e14 prophage are needed to test this hypothesis and directly measure the effects of chimeric *icd* on bacterial fitness, antibiotic tolerance, and persister phenotype formation^12,27^.

The conservative genetic changes observed in both protein alignments (Figures 1B and 2B) suggest that both chimeric proteins would be functional if they are indeed expressed. Supporting this interpretation, isocitrate dehydrogenase assays on cell lysates isolated before and after e14 excision by SOS induction indicate that the chimeric *icd* is both expressed and functional, albeit with lower activity than ancestral *icd*^26^. More generally, it is certainly possible that chimeric proteins formed by large genomic deletions could be highly beneficial, given the analogous finding that recombination between infectious lytic phage and a defunct prophage can dramatically improve phage fitness during coevolutionary phage training^33^.

Recent work by Kaul et al. has reported the evolution of novel gene fusions by large genomic deletions in both the LTEE and across the microbial tree of life^10^. In addition, uz-Zaman et al. has reported evidence of *de novo* gene birth in the LTEE^34^. This paper reports an additional mechanism that may contribute to novel genetic variation in microbial evolution experiments, and points to the need for quantitative estimates of the prevalence and importance of each mechanism for the evolution of novel genetic functions across the tree of life.

## Methods

### Genomic analysis of a 22-day fluoroquinolone-resistance evolution experiment

Genomic data published by Mohiuddin et al.^12^ were downloaded from ArrayExpress (https://www.ebi.ac.uk/biostudies/arrayexpress/studies/E-MTAB-14682, last accessed May 13, 2026). Also see Study ERP166963 (https://www.ebi.ac.uk/ena/browser/view/ERP166963, last accessed May 13, 2026) with Study Accession PRJEB83321 in the European Nucleotide Archive. *breseq* v0.37 was used to call mutations in these genomes using the *Escherichia coli* K-12 MG1655 reference genome in NCBI Refseq (GCF_000005845.2_ASM584v2). Mutations in the resequenced ancestral K-12 clone used for this experiment were applied to the NCBI RefSeq genome using the *gdtools* utility program distributed with *breseq* to generate an updated reference genome, which was then used to analyze mutations in each endpoint population. Resulting genomediff-format output files produced by *breseq* were collated using a Python 3.12 script called *process-gdiffs*.*py*, and statistical evidence for gene copy number amplifications was analyzed using an R 4.5.2 script called *copy-number-analysis*.*R*. Ancestral and evolved *icd* and *icdC* sequences were extracted from the K-12 reference genome and aligned using MAFFT^35^ v7.526 with a Python 3.12 script called *extract_and_align_icd_seqs*.*py*.

### Genomic analysis of manB-cpsG deletions in the LTEE

Mutations in evolved LTEE genomes were retrieved from https://barricklab.org/shiny/LTEE-Ecoli/ on May 4 2026. Ancestral and evolved *cpsG* homologs were extracted from the *E. coli* B str. REL606 reference genome and aligned using MAFFT^35^ v7.526 with a Python 3.12 script called *extract_and_align_cpsG_seqs*.*py*.

### Data visualization

The ancestral loci in the K-12 and REL606 genome before chimeric gene formation by interstitial deletions were visualized using Proksee^23^ and Benchling^22^. Visualizations of protein alignments were made using R 4.5.2 scripts called *plot_icd_alignment*.*R* and *plot_manB-cpsG_alignment*.*R*.

## Acknowledgements

We thank Nkrumah Grant, Jeffrey Barrick, Richard Lenski, and Sayed Golam Mohiuddin for helpful discussions. This work was partially funded by the New Jersey Agricultural Experiment Station (startup funds to R.M.) by the Rutgers Health BMIHAI Pilot Grant Program (R.M) and NIH/NIAID grant R01-AI143643 (M.O.) The authors declare that they have no potential conflicts of interest.

## SUPPLEMENTARY TABLES AND FIGURES

**Table S1.**
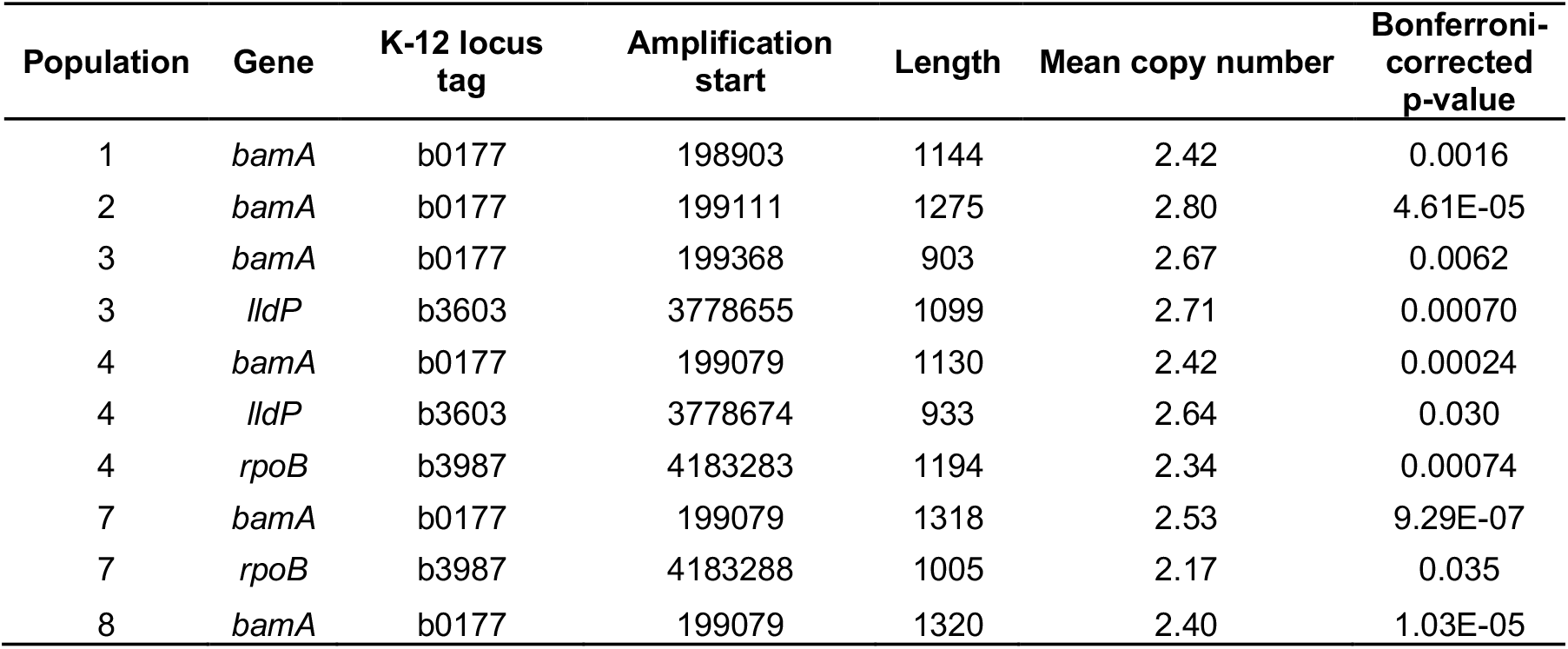
Gene amplifications in experiment reported by Mohiuddin et al. (2025).

| Population | Gene | K-12 locus tag | Amplification start | Length | Mean copy number | Bonferroni-corrected p-value |
| --- | --- | --- | --- | --- | --- | --- |
| 1 | *bamA* | b0177 | 198903 | 1144 | 2.42 | 0.0016 |
| 2 | *bamA* | b0177 | 199111 | 1275 | 2.80 | 4.61E-05 |
| 3 | *bamA* | b0177 | 199368 | 903 | 2.67 | 0.0062 |
| 3 | *lldP* | b3603 | 3778655 | 1099 | 2.71 | 0.00070 |
| 4 | *bamA* | b0177 | 199079 | 1130 | 2.42 | 0.00024 |
| 4 | *lldP* | b3603 | 3778674 | 933 | 2.64 | 0.030 |
| 4 | *rpoB* | b3987 | 4183283 | 1194 | 2.34 | 0.00074 |
| 7 | *bamA* | b0177 | 199079 | 1318 | 2.53 | 9.29E-07 |
| 7 | *rpoB* | b3987 | 4183288 | 1005 | 2.17 | 0.035 |
| 8 | *bamA* | b0177 | 199079 | 1320 | 2.40 | 1.03E-05 |

## REFERENCES

1 Conant, G. C. & Wagner, A. The rarity of gene shuffling in conserved genes. Genome biology 6, R50 (2005).

2 Xia, S., Chen, J., Arsala, D., Emerson, J. & Long, M. Functional innovation through new genes as a general evolutionary process. Nature genetics 57, 295–309 (2025).

3 Silhavy, T. J. Gene fusions. Journal of bacteriology 182, 5935–5938 (2000).

4 Ahsan, M. U., Liu, Q., Perdomo, J. E., Fang, L. & Wang, K. A survey of algorithms for the detection of genomic structural variants from long-read sequencing data. Nature methods 20, 1143–1158 (2023).

5 Chen, L. et al. Short-and long-read metagenomics expand individualized structural variations in gut microbiomes. Nature communications 13, 3175 (2022).

6 Rogers, R. L., Bedford, T. & Hartl, D. L. Formation and longevity of chimeric and duplicate genes in Drosophila melanogaster. Genetics 181, 313–322 (2009).

7 Smithers, B., Oates, M. & Gough, J. ‘Why genes in pieces?’—revisited. Nucleic Acids Research 47, 4970–4973 (2019).

8 Preska Steinberg, A. & Kussell, E. How recombination and clonal evolution shape bacterial lineages and genomes. Genetics 231, iyaf115 (2025).

9 Wolff, R. & Garud, N. R. Gene-specific selective sweeps are pervasive across human gut microbiomes. Nature 650, 710–717 (2026).

10 Kaul, A., Rossine, F., Břinda, K. & Baym, M. Novel genes arise from genomic deletions across the bacterial tree of life. bioRxiv (2026).

11 Panagopoulos, I. & Heim, S. Interstitial deletions generating fusion genes. Cancer Genomics & Proteomics 18, 167–196 (2021).

12 Mohiuddin, S. G., Kavousi, P., Figueroa, D., Ghosh, S. & Orman, M. A. The diverse phenotypic and mutational landscape induced by fluoroquinolone treatment. Msystems 10, e00713–00725 (2025).

13 Maddamsetti, R. et al. Duplicated antibiotic resistance genes reveal ongoing selection and horizontal gene transfer in bacteria. Nature Communications 15, 1449 (2024).

14 Levy, S. F. et al. Quantitative evolutionary dynamics using high-resolution lineage tracking. Nature 519, 181–186 (2015).

15 Schenk, M. F. et al. Population size mediates the contribution of high-rate and large-benefit mutations to parallel evolution. Nature Ecology & Evolution 6, 439–447 (2022).

16 Deatherage, D. E. & Barrick, J. E. in Engineering and analyzing multicellular systems: methods and protocols 165–188 (Springer, 2014).

17 Barrick, J. E. et al. Identifying structural variation in haploid microbial genomes from short-read resequencing data using breseq. BMC genomics 15, 1039 (2014).

18 Deatherage, D. E., Traverse, C. C., Wolf, L. N. & Barrick, J. E. Detecting rare structural variation in evolving microbial populations from new sequence junctions using breseq. Frontiers in genetics 5, 468 (2015).

19 Blount, Z. D. et al. Genomic and phenotypic evolution of Escherichia coli in a novel citrate-only resource environment. Elife 9, e55414 (2020).

20 Manuse, S. et al. Bacterial persisters are a stochastically formed subpopulation of low-energy cells. PLoS biology 19, e3001194 (2021).

21 Lopatkin, A. J. et al. Clinically relevant mutations in core metabolic genes confer antibiotic resistance. Science 371, eaba0862 (2021).

22 Benchling [Biology Software], < https://benchling.com. > (2026).

23 Grant, J. R. et al. Proksee: in-depth characterization and visualization of bacterial genomes. Nucleic acids research 51, W484–W492 (2023).

24 Greener, A. & Hill, C. Identification of a novel genetic element in Escherichia coli K-12. Journal of bacteriology 144, 312–321 (1980).

25 Brody, H. & Hill, C. Attachment site of the genetic element e14. Journal of bacteriology 170, 2040–2044 (1988).

26 Hill, C. W., Gray, J. A. & Brody, H. Use of the isocitrate dehydrogenase structural gene for attachment of e14 in Escherichia coli K-12. Journal of bacteriology 171, 4083–4084 (1989).

27 Sulaiman, J. E. & Lam, H. Proteomic investigation of tolerant Escherichia coli populations from cyclic antibiotic treatment. Journal of proteome research 19, 900–913 (2020).

28 Tenaillon, O. et al. Tempo and mode of genome evolution in a 50,000-generation experiment. Nature 536, 165–170 (2016).

29 Stevenson, G., Andrianopoulos, K., Hobbs, M. & Reeves, P. R. Organization of the Escherichia coli K-12 gene cluster responsible for production of the extracellular polysaccharide colanic acid. Journal of bacteriology 178, 4885–4893 (1996).

30 Schneider, D. et al. Genomic comparisons among Escherichia coli strains B, K-12, and O157:H7 using IS elements as molecular markers. BMC Microbiology 2, 18 (2002). 10.1186/1471-2180-2-18

31 Good, B. H., McDonald, M. J., Barrick, J. E., Lenski, R. E. & Desai, M. M. The dynamics of molecular evolution over 60,000 generations. Nature 551, 45–50 (2017).

32 Brody, H., Greener, A. & Hill, C. W. Excision and reintegration of the Escherichia coli K-12 chromosomal element e14. J Bacteriol 161, 1112–1117 (1985). 10.1128/jb.161.3.1112-1117.1985

33 Borin, J. M., Avrani, S., Barrick, J. E., Petrie, K. L. & Meyer, J. R. Coevolutionary phage training leads to greater bacterial suppression and delays the evolution of phage resistance. Proceedings of the National Academy of Sciences 118, e2104592118 (2021).

34 Uz-Zaman, M. H., D’Alton, S., Barrick, J. E. & Ochman, H. Promoter recruitment drives the emergence of proto-genes in a long-term evolution experiment with Escherichia coli. Plos Biology 22, e3002418 (2024).

35 Katoh, K. & Standley, D. M. MAFFT multiple sequence alignment software version 7: improvements in performance and usability. Molecular biology and evolution 30, 772–780 (2013).

